# Cell type-specific differences in redox regulation and proliferation after low UVA doses

**DOI:** 10.1101/425793

**Authors:** Sylwia Ciesielska, Patryk Bil, Karolina Gajda, Aleksandra Poterala-Hejmo, Dorota Hudy, Joanna Rzeszowska-Wolny

## Abstract

Ultraviolet A (UVA) radiation is harmful for living organisms but in low doses may stimulate cell proliferation. Our aim was to examine the relationships between exposure to different low UVA doses, cellular proliferation, and changes in cellular reactive oxygen species levels. In human colon cancer (HCT116) and melanoma (Me45) cells exposed to UVA doses comparable to environmental, the highest doses (30-50 kJ/m^2^) reduced clonogenic potential but some lower doses (1 and 10 kJ/m^2^) induced proliferation. This effect was cell type and dose specific. In both cell lines the levels of reactive oxygen species and nitric oxide fluctuated with dynamics which were influenced differently by UVA; in Me45 cells decreased proliferation accompanied the changes in the dynamics of H_2_O_2_ while in HCT116 cells those of superoxide. Genes coding for proteins engaged in redox systems were expressed differently in each cell line; transcripts for thioredoxin, peroxiredoxin and glutathione peroxidase showed higher expression in HCT116 cells whereas those for glutathione transferases and copper chaperone were more abundant in Me45 cells. We conclude that these two cell types utilize different pathways for regulating their redox status. Many mechanisms engaged in maintaining cellular redox balance have been described. Here we show that the different cellular responses to a stimulus such as a specific dose of UVA may be consequences of the use of different redox control pathways. Assays of superoxide and hydrogen peroxide level changes after exposure to UVA may clarify mechanisms of cellular redox regulation and help in understanding responses to stressing factors.

## Introduction

Ultraviolet radiation is the non-ionizing part of the electromagnetic radiation spectrum with a wavelength of 100-400 nm, invisible to human sight. The sun is a natural emitter of UV divided into three main fractions UVA (315-400 nm), UVB (280-315 nm), and UVC (100-280 nm), but most of this radiation is blocked by the atmosphere (1,2). UVA constitutes the largest part (~95%) of UV radiation that reaches the Earth’s surface (3), whereas UVB represents only 4-5% (1). In irradiated humans UVA reaches the dermis and hypodermis and has no direct impact on DNA, but it can influence cellular structures indirectly by induction of reactive oxygen species (ROS) which can damage macromolecules (4,1). For a long time UV was regarded as damaging for cells and organisms (5), but since a few decades it is known that low doses can also stimulate proliferation of cells; however, the mechanisms underlying this phenomenon are not completely understood (3,1,6,7).

Studies of signaling pathways in conditions where UVA stimulates cell proliferation show changes in the levels of proteins engaged in controlling proliferation such as cyclin D1 (8,9), Pin1 (3), and Kin17 (10) or activation of epidermal growth factor receptor (EGFR) which is strongly mitogenic in many cell types (8). Experiments on mice showed that UVA can accelerate tumor growth (2,11).

One effect of exposure to UV is induction of ROS in cells, including different reactive molecules and free radicals derived from molecular oxygen (12) which together with reactive nitrogen species (RNS) play important roles in regulation of cell signaling and survival (reviewed in 13). ROS can exert opposing effects, inducing cell damage and death or stimulating proliferation by protein modifications and participation in signaling pathways (14-23). Many complex mechanisms guard redox homeostasis, the balance between generation and elimination of ROS and antioxidant systems, such as superoxide dismutase, catalase or glutathione peroxidases which participate in these control systems (24,22). The role of ROS in stimulating proliferation by low doses of UVA was supported by experiments in which irradiation with a low-power diode laser increased ROS production accompanied by increased cell proliferation which was prevented by addition of catalase or superoxide dismutase (9), suggesting that ROS are at least partly involved in stimulating proliferation (19). ROS in cells originate both from external sources and as byproducts of cellular processes (24, 9, 20, 21). Low levels of ROS stimulate cell proliferation by activating signaling pathways connected with growth factors, causing increased cell cycle progression, while higher levels show toxic effects causing cell death or senescence (24, 25). RNS include nitric oxide (NO), a highly reactive gas synthesized from L-arginine by members of the nitric oxide synthase (NOS) family (26). NO modulates many cellular functions (27) by acting as a messenger for paracrine and autocrine communication and its production and degradation are strictly controlled in different cell types (28). All cells of multicellular organisms produce superoxide and NO, which appear to be the main radicals responsible for the regulation of cellular redox homeostasis. This regulation is especially important in the presence of external ROS sources, because cells do not distinguish between endogenously- and exogenously-generated ROS. The main endogenous sources of superoxide are electron leakage from the mitochondrial respiratory chain and NADPH oxidases (NOXs), a family of enzymes dedicated to the production of ROS in a variety of cells and tissues (reviewed in 29, 20, 30). The generation of superoxide is highly conserved across all eukaryotic life and is strictly regulated by antioxidant enzymes and reducing agents (13,29), and the fluctuating level of ROS in cells has been postulated to be an important mechanism regulating progression through the cell cycle (31, 20, 22, 32).

As ROS and NO play an important role in many intra- and inter-cellular signaling pathways, participate in regulation of the cell cycle (reviewed in 20), and show increased levels after UV radiation (4) we have studied if and how changes in their levels in irradiated cells could be related to the effects of UVA on proliferation, using human melanoma (Me45) and colon cancer (HCT116) cells irradiated with UVA. We show that some low doses, specific for each cell line, stimulate clonogenic survival whereas other, even lower doses inhibit proliferation. Comparison of the changes in the intracellular levels of ROS, NO, and superoxide (O_2_^-^) after irradiation with stimulating, suppressing, or neutral UVA doses suggests that these cell lines regulate their ROS levels by different pathways, and that it is the dynamics of superoxide or H_2_O_2_ levels which plays a crucial role in growth stimulation or inhibition.

## Materials and methods

### Cell lines and culture

Human melanoma cells (Me45, established in the Center of Oncology in Gliwice from a lymph node metastasis of skin melanoma; 33) and human colorectal carcinoma cells (HCT116; p53+/+, ATCC) were maintained in DMEM/F12 medium (PAN Biotech. Aidenbach, Germany, cat, #P04-41150) enriched with 10% fetal bovine serum (EURx, Gdansk, Poland cat# E5050-03-500) at 37 °C in a humidified atmosphere enriched in 5% CO_2_. The cells, 1000-5000 per dish, were irradiated at room temperature (21°C) in culture plates (Sarstedt, Numbrecht, Germany cat# 83.3900) (covers opened) with various doses (0.05–50 kJ/m^2^) of UVA (365 nm) generated by a UV crosslinker (model CL-1000, UVP, Upland, CA, USA) and used for clonogenic survival assays.

### Clonogenic survival asssays

Control and irradiated cells were seeded in 60-mm dishes at 1000-5000 cells/dish and incubated from 5 to 14 days (depending on the cell line) at 37°C in a humidified atmosphere. The colonies were fixed with 2 ml cold 96% ethanol for 3 min, than washed with PBS (PAN Biotech., Aidenbach, Germany, cat. no. P04-36500) and stained with 0.5% methylene blue in 50% ethanol. Cells in colonies containing more than 50 cells (estimated under the microscope) were counted and the surviving fraction was calculated as the plating efficiency of irradiated cells relative to that of control un-irradiated cells.

### Intracellular reactive oxygen species levels

To quantitate intracellular ROS, 100.000 cells were seeded, growing cells were collected by trypsinization, suspended in culture medium to which 2′,7′-dichlorofluorescein diacetate (DCFH-DA; Sigma-Aldrich, St. Louis, USA, Cat#287810) was added to final concentration of 30µM. Cells were incubated for 30 min at 37°C in the dark, washed with medium, suspended in PBS, and kept for 15 min on ice in the dark. Fluorescence was measured by flow cytometry (Becton Dickinson FACS Canto) using the FITC configuration (488 nm laser line, LP mirror 503, BP filter 530/30), usually 10,000 cells were assayed per sample. To assess superoxide radicals in living cells, MitoSox Red fluorogenic reagent (Thermo Fisher Scientific, Waltham, USA, cat. no. M36008) was used (34, 35). Cells were collected, suspended in PBS (20,000 cells/300µl), incubated with MitoSox Red (5 µM final concentration) for 20 min at 37°C in the dark, and washed and resuspended in PBS. Samples were kept on ice until analysis by flow cytometry (Becton Dickinson FACS Canto, 488 nm laser line, LP mirror 566, BP filter 585/42), measuring 10,000 cell per sample. To assess NO, cells were incubated with 1µM 4-amino-5-methylamino-2′,7′-difluorescein diacetate (DAF-FM, Thermo Fisher Scientific, Waltham, USA, cat.# D23844) for 30 min in dark conditions at 37°C and washed with PBS. The fluorescence intensity of 10,000 cells was measured by flow cytometry using the FITC configuration (488 nm laser line. LP mirror 503, BP filter 530/30).

Results are expressed as mean fluorescence intensities ±SD from three independent experiments.

### Fluorescence microscopy and image analysis

Fluorescent microscopy assays of superoxide and NO were performed with the same fluorescent reagents as for cytometry (MitoSOX Red and DAF-FM diacetate, Thermo Fisher Scientific, Waltham, USA). HCT116 and Me45 cells were seeded at 10,000 cells per well in 4-well cell culture chambers (Sarstedt, Numbrecht, Germany, cat# 94.6140.402), grown in DMEM medium supplemented with 10% fetal bovine serum for 24 hours at 37°C in standard conditions, and labelled with MitoSOX Red (2.5µM) in the first well, DAF-FM Diacetate (2.5µM) in the second well, both dyes in the third well, and no dye in the last (control) well. Cells were incubated for 20 minutes at 37°C in a humidified atmosphere enriched with 5% CO_2_, the culture medium was removed, the cells were washed with PBS, fixed with 0.5 ml of cold 70% ethanol per well for 10 minutes, and washed with the same volume of deionized water for 3 minutes. Slides with fixed cells were covered with mounting gel and a cover glass. Images were captured with an Olympus BX43 microscope with a 40x objective and a CoolLED precisExcite fluorescence excitation system. Red and green fluorescence and transparent light images were obtained for 10 areas containing cells stained with both fluorescent dyes on each slide and analyzed with Matlab 2016b software using the functions corrcoef() and scatter() to detect correlation between the values of corresponding pixels in both fluorescence images.

### Expression of genes coding for proteins engaged in cellular redox processes

We identified 574 genes which are directly or indirectly engaged in redox processes, using GO terms such as oxide, superoxide, nitric oxide, hydrogen peroxide, ROS and reactive oxygen species. The levels of transcripts of these genes in non-irradiated HCT116 and Me45 cells were extracted from our earlier Affymetrix microarray experiments (32, 17) whose results are available in the ArrayExpress database under accession number E-MEXP-2623. All data are MIAME compliant.

### Assay of total and oxidized glutathione levels

For assays of total glutathione we used Rahman et al.’s modification (36) of the colorimetric assay originally proposed by Vandeputte et al. (37) which is based on the reaction of GSH with 5,5′-dithio-bis (2-nitrobenzoic acid) (DTNB, Singma-Aldrich,Saint Louis, USA, cat# D-8130) which produces 5-thio-2-nitrobenzoic acid (TNB) and its adduct with oxidized glutathione (GS-TNB). The disulfide product was reduced by glutathione reductase (0.2 U) (Sigma-Aldrich, Saint Louis, USA, cat. no. G-3664)) in the presence of 0.8mM NADPH (Sigma-Aldrich, Saint Louis, USA, cat. no. D-8130). The TNB chromophore was measured at 412 nm in a microplate (96-plate) reader (Epoch, Biotek, Winooski, USA). For measurements of oxidized glutathione (GSSG) levels, cell extracts made by sonication in 0.1% Triton X-100 (Sigma-Aldrich, Saint Louis, USA, cat# T8787) and 0.6% sulfosalicylic acid (Sigma-Aldrich, Saint Louis, USA, cat# S-2130) in 0.05M potassium phosphate buffer pH 7.2 containing 1 mM EDTA (KPE) buffer were treated with 2-vinylpyridine (Sigma-Aldrich, Saint Louis, USA, cat# 132292) for 1 h at room temperature, excess 2-vinylpyridine was neutralized with triethanolamine (Sigma-Aldrich, Saint Louis, USA, cat# T1377), and the enzymatic recycling and reaction with DTNB was carried as described above.

### Statistical analyses

At least three replicates of all experiments were performed and results are expressed as means ± SD and summarized as percentages relative to the appropriate controls. Differences between samples were regarded as statistically significant at a p-value < 0.05 calculated by the two-sided Student t-test. Correlations between time course changes in irradiated and control cells were calculated using Pearson’s test and are presented as correlation coefficients.

## Results

### UVA induced proliferation changes are dose and cell-type specific

HCT116 and Me45 cells were exposed to a range of UVA radiation doses (0.05, 0.1, 0.25, 0.5, 1, 5, 10, 15, 20, 30, 40, or 50 kJ/m^2^) and their proliferation was studied by clonogenic tests. Some doses stimulated proliferation and others suppressed proliferation when compared to un-irradiated controls in both cell lines, although they responded differently and the doses that increased clonogenicity were specific for each cell line (**Fig. 1**). HCT116 cells showed a statistically significant increase of colony formation after exposure to 10 kJ/m^2^ (p-value 0.02) and a decrease after 0.1, 40, and 50 kJ/m^2^ (p-values 0.02, 0.05 and <0.01). The clonogenicity of Me45 cells increased after irradiation with 1 and 10 kJ/m^2^ (p-value <0.01) but was reduced after 15 to 50 kJ/m^2^ (p-values 0.01, 0.01, 0.045, 0.04 and <0.01 respectively).

**Fig 1.**
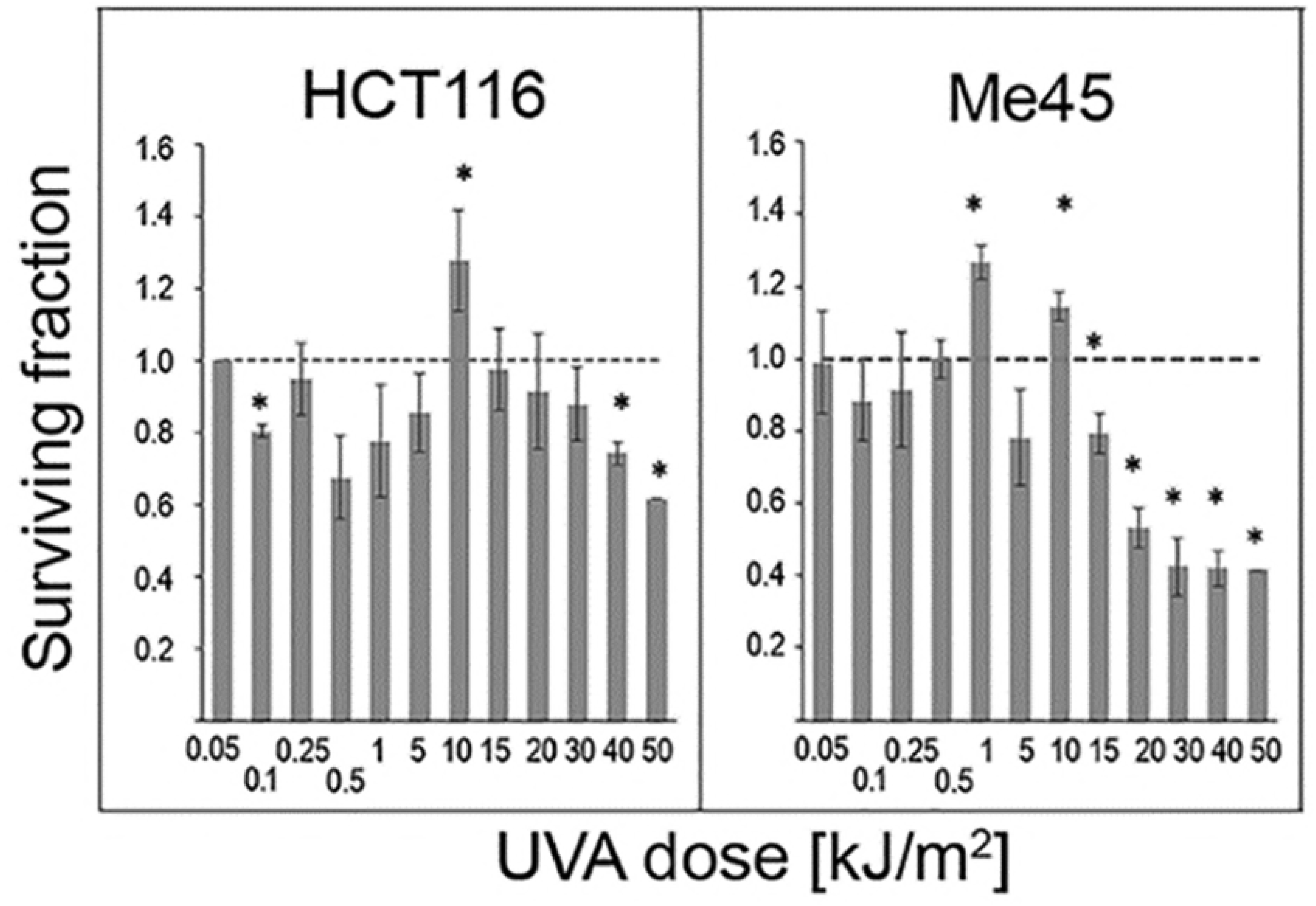
Clonogenicity of human cells after exposure to different UVA doses. (A) HCT116 cells, (B) Me45 cells. Data show the mean and SD of 3 experiments. Asterisks denote statistical significance of differences between irradiated and control samples with a p-value <0.05. The horizontal dashed line represents the control level.

### Low UVA doses do not significantly influence average levels of ROS

We used specific fluorescent probes and flow cytometry to compare the levels of ROS and NO in cells irradiated with different UVA doses with those in control cells. **Fig 2** shows the effect of UVA on the level of superoxide detected by MitoSox, of NO detected by DAF-FM, and of ROS detected by DCFH-DA. The average values for each dose were calculated from all twelve assays performed in different experiments and at different time points.

**Fig 2.**
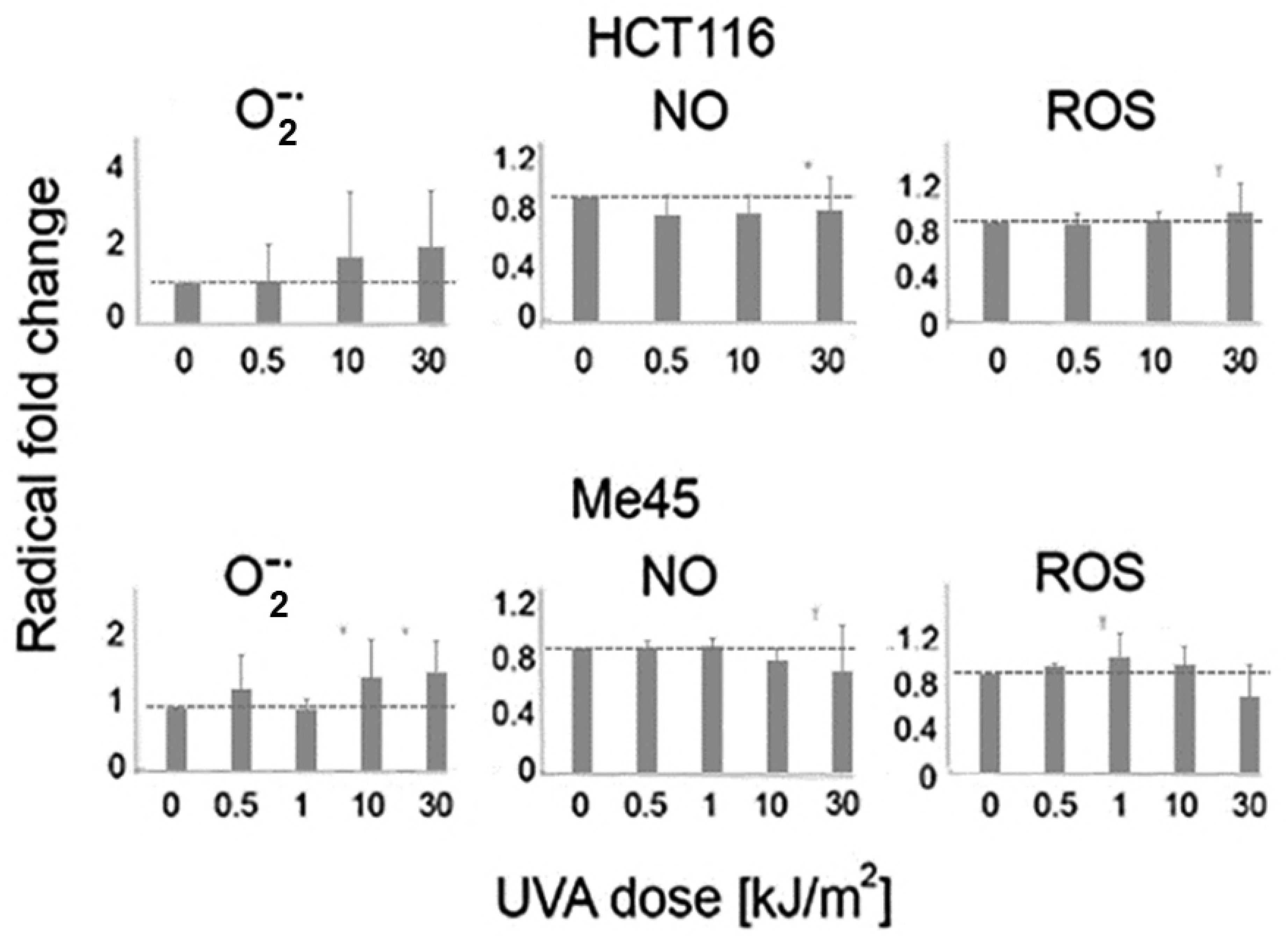
Average levels of ROS and NO in HCT116 and Me45 do not significantly change after exposure of cells to different UVA doses. The levels were measured four times during 24 h in control and cells irradiated with different UVA doses and experiment was repeated 4 times. The results are presented as fold change in irradiated cells versus non-irradiated controls. Data show the mean and SD of 4 experiments.

Average superoxide levels showed a tendency to increase with higher UVA dose in both cell lines, but the increases were not statistically significant. NO levels did not change or decreased slightly with higher doses. The levels of ROS detected with DCFH-DA also did not change in irradiated HCT116 cells, but Me45 cells showed small irregular increases with lower doses and decreases with higher doses. This probe detects several different radicals and was first used for detection of H_2_O_2_ (38, 39), and it seems probable that the ROS changes detected by this probe mainly reflect changes of H_2_O_2_ levels. None of the differences in average levels of ROS or NO radicals between control and irradiated cells were statistically significant.

### ROS level dynamics change differently after different UVA doses

Although the UVA doses which we used did not change the average ROS levels significantly, they influenced fluctuations of these levels. The time course changes of the levels of ROS assayed by DCFH-DA, of superoxide, and of NO in cells irradiated with a particular dose or not irradiated are shown in **Fig 3**. Me45 and HCT116 cells responded to different doses with very different kinetics of radical levels and these dynamics of changes were cell type-specific. At first sight it is difficult to identify features which could be correlated with the increased or decreased clonogenic potential observed after irradiation with some doses.

**Fig 3.**
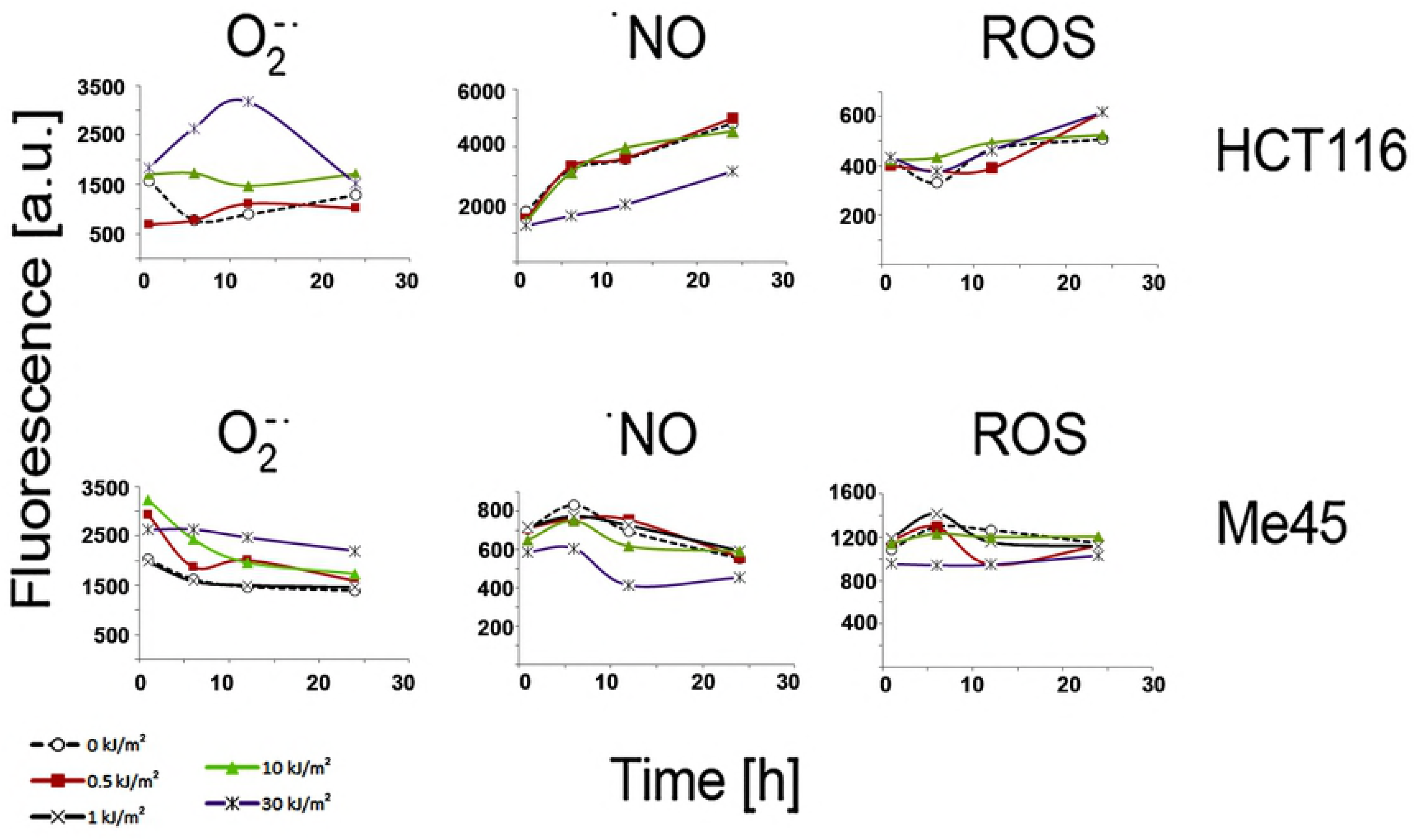
The dynamics of the levels of superoxide, nitric oxide and ROS detected by DCFH-DA in control and UVA irradiated cells. Each curve represents the results after exposure to a particular UV dose shown on the right; data are means from three experiments and error bars are not shown for clarity.

To evaluate the similarity between radical dynamics in UVA-irradiated and control cells, we calculated correlation coefficients using Pearson’s test. The dynamics of NO levels did not change significantly after exposure of cells to any of the UVA doses studied, and the increases and decreases appeared at similar time points in control and irradiated cells. The correlation coefficients between cells irradiated with different doses or not irradiated were >0.9 for HCT116 cells and >0.8 for three out of four doses in Me45 cells (**Table 1**). This positive correlation suggests that the changes of NO levels are strictly controlled in both cell lines after both stimulating or inhibiting proliferation UVA doses.

**Table 1.** Correlation coefficients for the degree of similarity between radical dynamics in UV-irradiated vs. control cells.

|  |  |  |  |
| --- | --- | --- | --- |
|  | **Superoxide<sup>1</sup>** | **ROS<sup>2</sup>** | **NO<sup>3</sup>** |
| <u>HCT116 cells</u> |  |  |  |
| 0.5 kJ/m<sup>2</sup> | **-0.38** | 0.7 | 1* |
| 10 kJ/m<sup>2</sup> | 0.37 | 0.81 | 0.97* |
| 30 kJ/m<sup>2</sup> | **-0.79** | 0.88 | 0.94* |
| <u>Me45 cells</u> |  |  |  |
| 0.5 kJ/m<sup>2</sup> | 0.95* | **-0.01** | 0.89* |
| 1 kJ/m<sup>2</sup> | 0.99* | 0.59 | 0.89* |
| 10 kJ/m<sup>2</sup> | 0.99* | 0.83 | 0.93* |
| 30 kJ/m<sup>2</sup> | 0.98* | **-0.47** | 0.67 |
| <sup>1</sup>measured by MitoSox. <sup>2</sup>detected by DCFH-DA (mainly H<sub>2</sub>O<sub>2</sub>). <sup>3</sup>measured by DAF-FM; *Pearson's correlation p-value <0.05. Bold values indicate changes from a positive to a negative correlation coefficient. |  |  |  |

The superoxide level dynamics in Me45 cells irradiated with any dose were highly correlated with those in control cells (**Table 1**). In contrast, in HCT116 cells this level showed clear differences between the effects of UVA doses which stimulated or did not stimulate clonogenic potential; the dynamics of superoxide levels after doses inhibiting proliferation were inversely correlated with those in control cells, while after doses which stimulated proliferation these levels were positively correlated with those in control cells; however the correlation coefficients were rather low. The dynamics of the level of ROS in Me45 cells assayed by DCFH-DA changed after irradiation in a manner similar to those of superoxide in HCT116 cells, proliferation-inhibiting doses showing a negative correlation and proliferation-stimulating doses a positive correlation with the dynamics in control cells.

### HCT116 and Me45 cells differ in intracellular level and localization of NO and superoxide

Me45 cells had a ~5 times lower level of NO than HCT116 cells but higher levels of ROS detected by DCFH and of superoxide, as assayed by flow cytometry (**Fig 4A**). Analysis of single cells using fluorescence microscopy showed that in both cell types, most NO and superoxide were co-localized as shown by a high positive correlation of their signals in single pixels. Rare HCT116 cells contained larger regions with a high NO and a low superoxide signal (for example, Fig. 4B) but similar regions were not seen in Me45 cells. Co-localization was significantly higher in Me45 than in HCT116 cells; Pearson’s correlation coefficients for all pixels in 10 fields containing 5 to 10 cells were 0.9 and 0.6 in Me45 and HCT116 cells, respectively.

**Fig 4.**
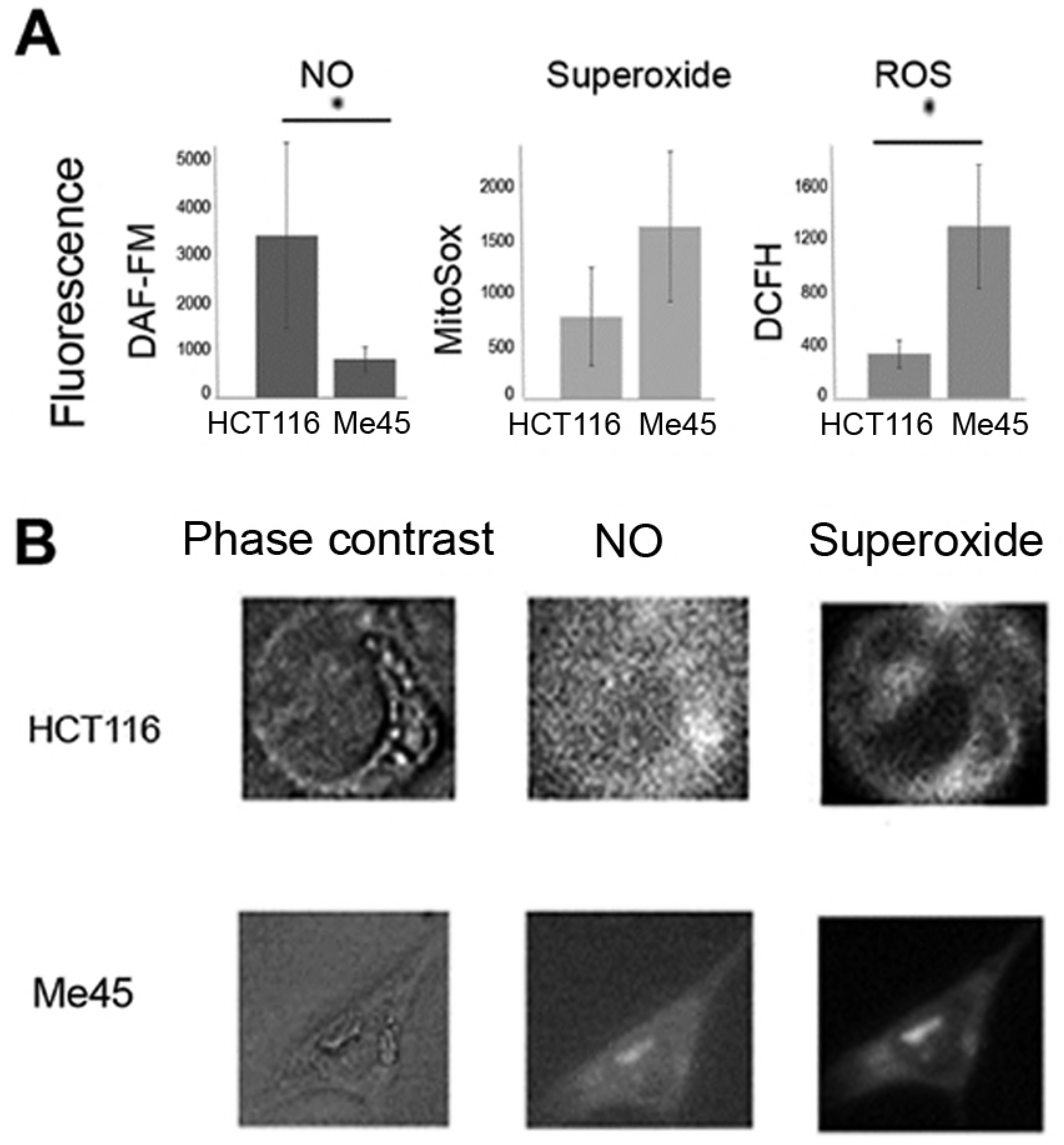
Nitric oxide and ROS in HCT116 and Me45 cells. A; mean levels of NO, superoxide and ROS detected by DCFH-DA measured in whole population of unirradiated cells by flow cytometry (average from 3 experiments), B; examples of superoxide and NO distribution in single HCT116 and Me45 cells observed by fluorescence microscopy, NO detected by fluorescence of DAF-FM diacetate and superoxide by MitoSOX Red.

### HCT116 and Me45 cells have different levels of some transcripts participating in redox systems

The differences in response to UVA and in radical levels in the two cell lines suggested that they use different mechanisms for the regulation of their redox status. To get more information on these mechanisms, we compared the expression of different genes coding for proteins engaged directly or indirectly in redox processes in each cell line. The expression levels of more than 500 candidate genes found on the basis of ontology terms were compared using our earlier microarray data for Me45 and HCT116 cells (17). The full list of these genes and their expression levels are given in Table S1 of the Supplement. Both cell lines express many genes engaged in redox regulation and expression of some of these genes is significantly higher in Me45 or HCT116 cells (Tables 2 and 3).

**Table 2.** Genes with higher expression in Me45 than in HCT116 cells.

| **Table 2. Genes with higher expression in Me45 than in HCT116 cells** |  |  |  |
| --- | --- | --- | --- |
| **Gene** | **Gene symbol** | **Transcript level [a.u.]<sup>1</sup>** | **Enrichment<sup>2</sup>** |
| Glutathione S-Transferase Mu 3 | *GSTM3* | 299 | 27.0 |
| Antioxidant 1 Copper Chaperone | *ATOX1* | 3336 | 11.3 |
| Thioredoxin Interacting Protein | *TXNIP* | 346 | 10.7 |
| Glutathione S-Transferase Alpha 4 | *GSTA4* | 316 | 3.0 |
| Peroxidasin | *PXDN* | 80 | 2.8 |
| Glutathione S-Transferase Kappa 1 | *GSTK1* | 734 | 2.4 |
| SH3 Domain Binding Glutamate Rich Protein Like 3 | *SH3BGRL3* | 552 | 2.3 |
| Cyclin Dependent Kinase 5 | *CDK5* | 565 | 2.2 |
| Cyclin Dependent Kinase 2 | *CDK2* | 359 | 2 |
| Glutathione S-Transferase Pi 1 | *GSTP1* | 1042 | 1.8 |
| Catalase | *CAT* | 402 | 1.6 |
| Glutathione S-Transferase Omega 1 | *GSTO1* | 2037 | 1.4 |
| Microsomal Glutathione S-Transferase 3 | *MGST3* | 1334 | 1.2 |
| Thioredoxin Reductase 1 | *TXNRD1* | 758.2 | 1.1 |
| <sup>1</sup>arbitrary units reflect normalized data from microarray experiment. <sup>2</sup>Fold change |  |  |  |

**Table 3.**
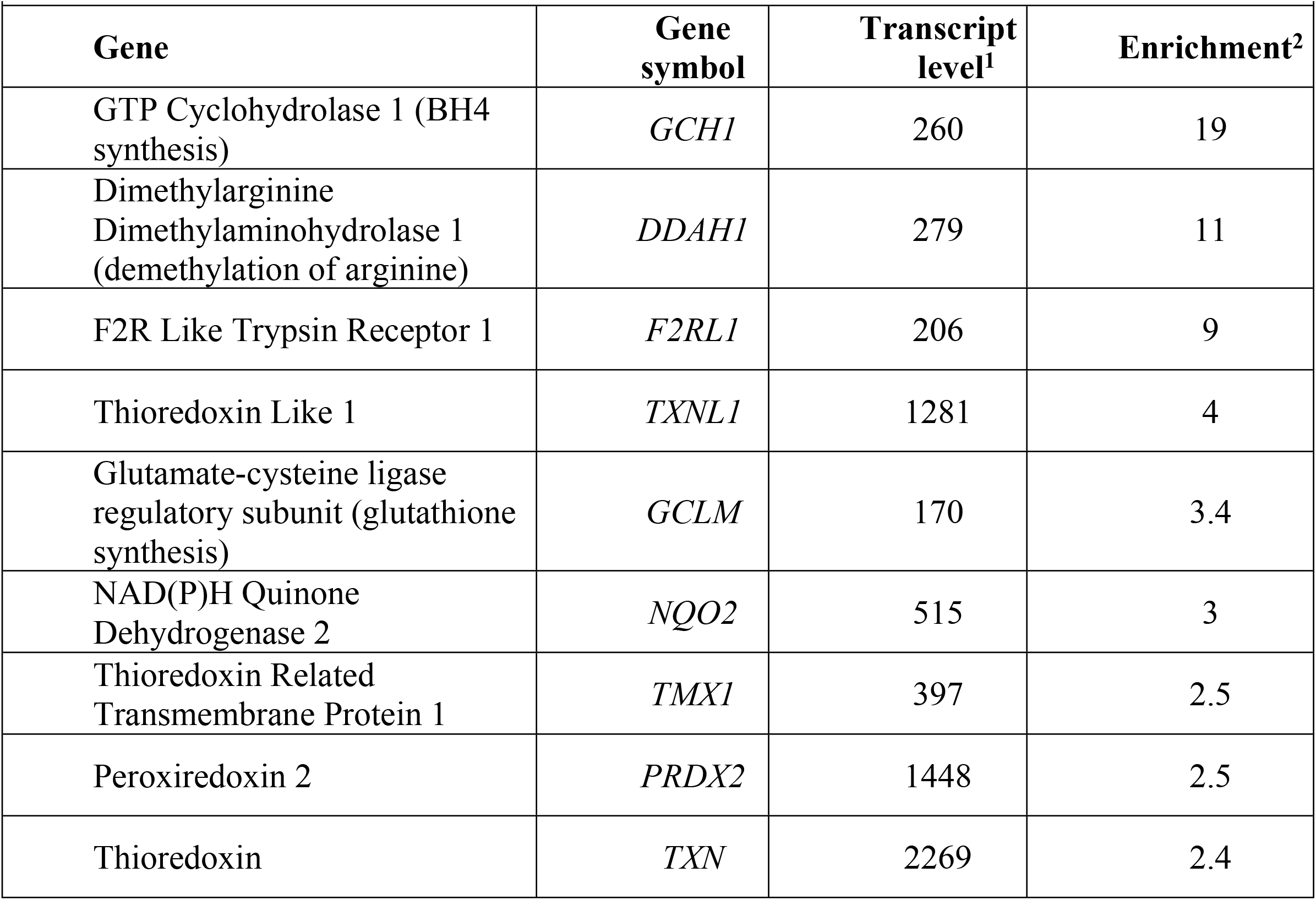
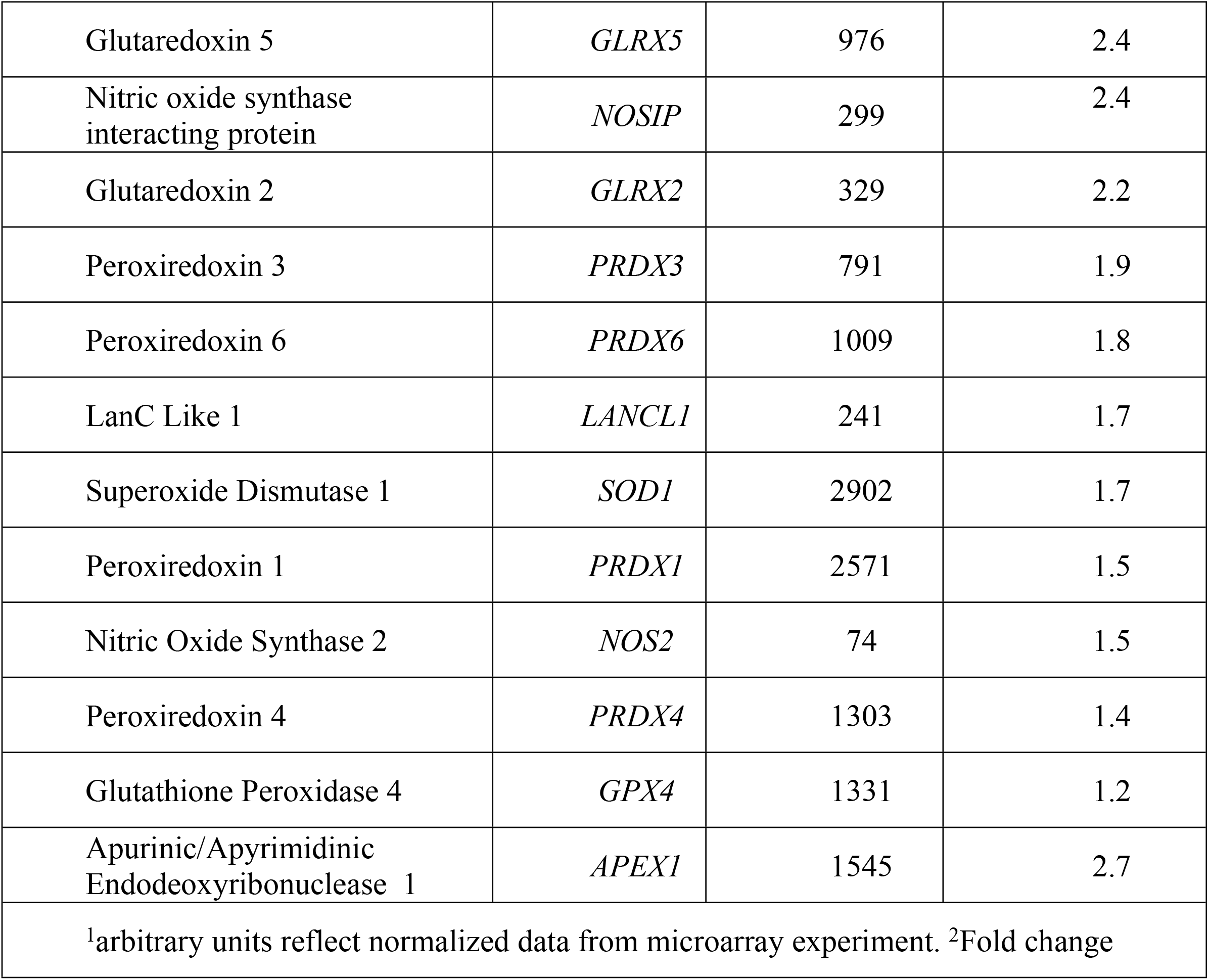
Genes with higher expression in HCT116 than in Me45 cells.

Me45 cells contain lower levels of transcripts for thioredoxin (TXN) and peroxyredoxin (PRDX) and higher levels of transcripts for thioredoxin-inhibiting protein (TXNIP). On the other hand, genes coding for glutathione S-transferases (GST) show higher expression in Me45 cells, with the GSTM3 transcript showing the largest difference. Transcripts for the antioxidant ATOX1, a copper chaperone which may increase activity of the protein SOD1 by providing copper ions and influence SOD3 gene expression as a transcription factor (40, 41), are more than 10 times more abundant in Me45 than in HCT116 cells. There are also some genes which are significantly more highly expressed in HCT116 cells, for example GTP cyclohydrolase 1 which codes for the first and rate-limiting enzyme in biosynthesis of tetrahydrobiopterin (BH4), a cofactor required for activity of nitric oxide synthases (42,43).

### Glutathione in HCT116 and Me45 cells

Glutathione is an important player in cell redox regulation (36) and the gene *GCLM* which codes for glutamate-cysteine ligase regulatory subunit, required for synthesis of glutathione, is more highly expressed in HCT116 than in Me45 cells (Table 3). We therefore compared the levels of reduced (GSH) and oxidized (GSSG) glutathione in these cells. The levels of total glutathione, GSH (~96% of the total), and of GSSG were lower in Me45 cells, but the differences were not statistically significant (**Fig 5**).

**Fig 5.**
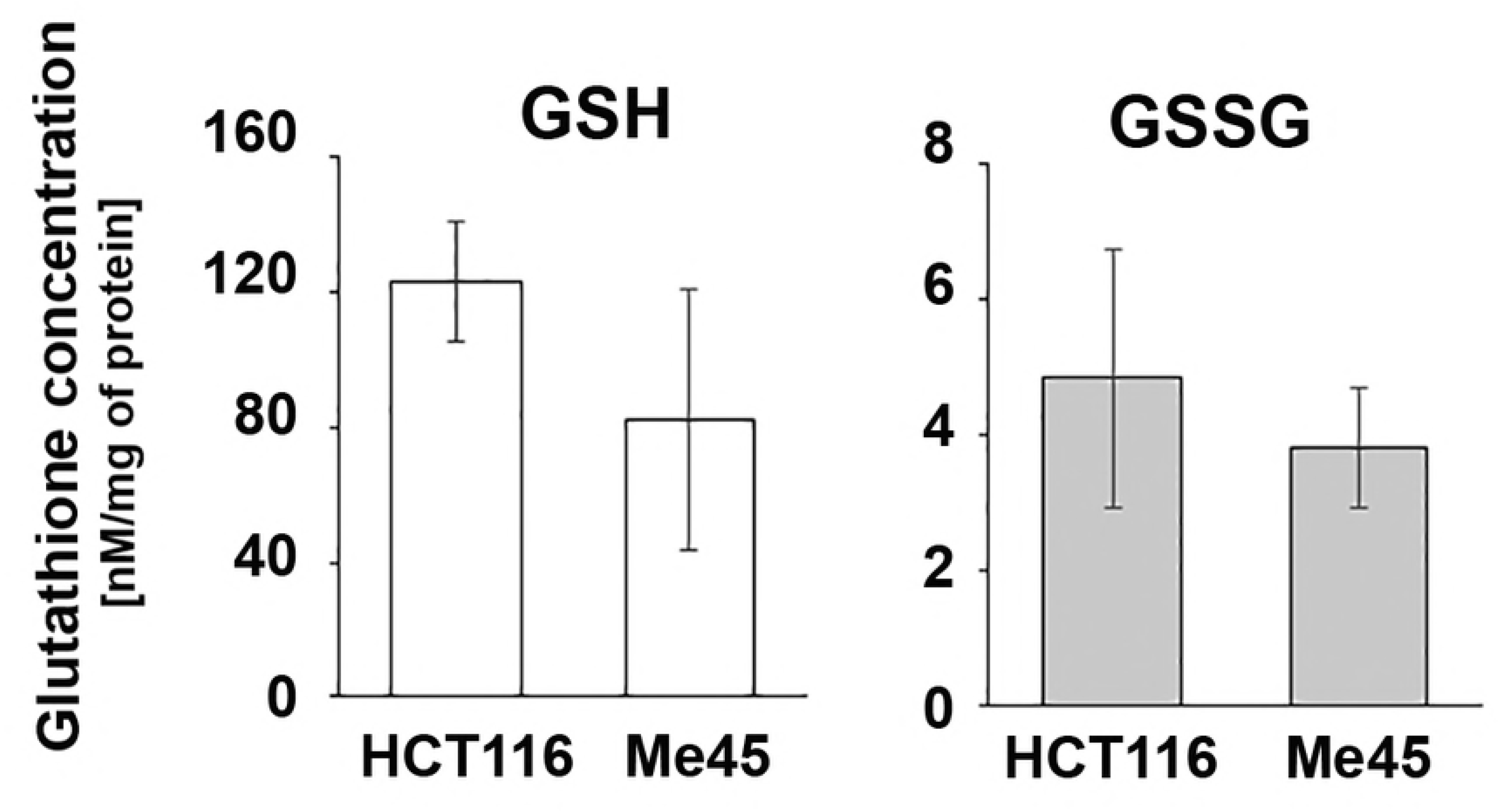
Levels of reduced (GSH) and oxidized (GSSG) glutathione in HCT116 and Me45 cells. Data show the mean and SD of 3 independent experiments.

## Discussion

### Stimulation of proliferation by UVA and fluctuations of intracellular ROS level

Stimulation of cell proliferation by UVA radiation at doses of 3-9 kJ/m^2^ has been known for a few decades. (9, 3). Here we show that doses in this range, but not exceeding 10 kJ/m^2^, increase the clonogenic potential of HCT116 and Me45 cells and that this effect is dose- and cell type-specific (Fig. 1). We further relate this specificity to cellular redox regulation, supporting a role for redox conditions and superoxide and NO in regulation of proliferation which was suggested 30 years ago (44, 14, 45 reviewed in 46,47). We hypothesized that the induction of cell proliferation by UVA may be caused by changes in intracellular levels of ROS and RNS (34, 35).

The levels of intracellular ROS, superoxide, and NO, assayed using specific probes, changed in time (Fig 3) in agreement with the fluctuations of ROS level observed by others and proposed to be important in regulation of the cell cycle (reviewed in 20). In some cases the kinetics of the changes of level after irradiation were highly correlated with those in control cells (Table 1); for example, in both cell types the general pattern of NO level change did not vary after irradiation although their levels differed (Fig 3), suggesting that the pattern of NO level change is important for regulatory mechanisms in both cell types. For other radicals, the correlation between irradiated and control cells was much lower and sometimes changed sign; for example, in Me45 cells the fluctuations of superoxide level did not vary after irradiation and were highly correlated with those in control cells, whereas in contrast the fluctuations in HCT116 cells varied depending on the UVA dose and were inversely correlated with those in control cells after proliferation-inhibiting doses, but were positively correlated after proliferation-stimulating doses. In HCT116 cells the DCFH-DA-detected ROS level changed more regularly than that in Me45 cells, while in Me45 cells irradiated with proliferation-inhibiting UVA doses it became inversely correlated compared to the dynamics in control cells. An increase of proliferation rate after irradiation was observed only if the fluctuations of ROS level retained their pattern in control cells, although conservation of the pattern of fluctuations of different radicals in both cell lines were important (Table 1). Overall, these results suggest that it is the pattern of fluctuations of radical levels, rather than the levels themselves, which influences proliferation rate after UVA irradiation and that each cell type may use different pathways to regulate cellular redox status.

### ROS-regulating pathways and their choice in HCT116 and Me45 cells

ROS participate in many signaling pathways, including those regulating the cell cycle and proliferation (24, 9, 20, 22), and their intracellular levels must be precisely controlled. The main players in regulation of cellular redox status are superoxide and NO which are produced by cells and interact with each other and with many other cellular molecules. Their levels are regulated by a series of feedback circuits, mainly based on peroxiredoxins, thioredoxins, glutathione, thioredoxin and glutathione reductases, NADPH, and enzymes engaged in production of superoxide or NO (reviewed in 48,49,50) (**Fig 6**). Fig 6 shows some proteins whose differential expression in Me45 and HCT116 cells may influence these pathways. Many other possible interactions of superoxide and ONOO^-^ occur, with themselves, with other proteins, CO_2_, antioxidants, and other compounds which result in creation of new radicals and interaction circuits which further influence the redox state of the cell and create additional regulatory sub-circuits, described in detail in many recent and older reviews (48,51,50,52). Nevertheless, the ROS regulatory circuits in Fig 6 seem to create the basic pathways for redox regulation in cells which may determine the character of radical level fluctuations.

**Fig 6.**
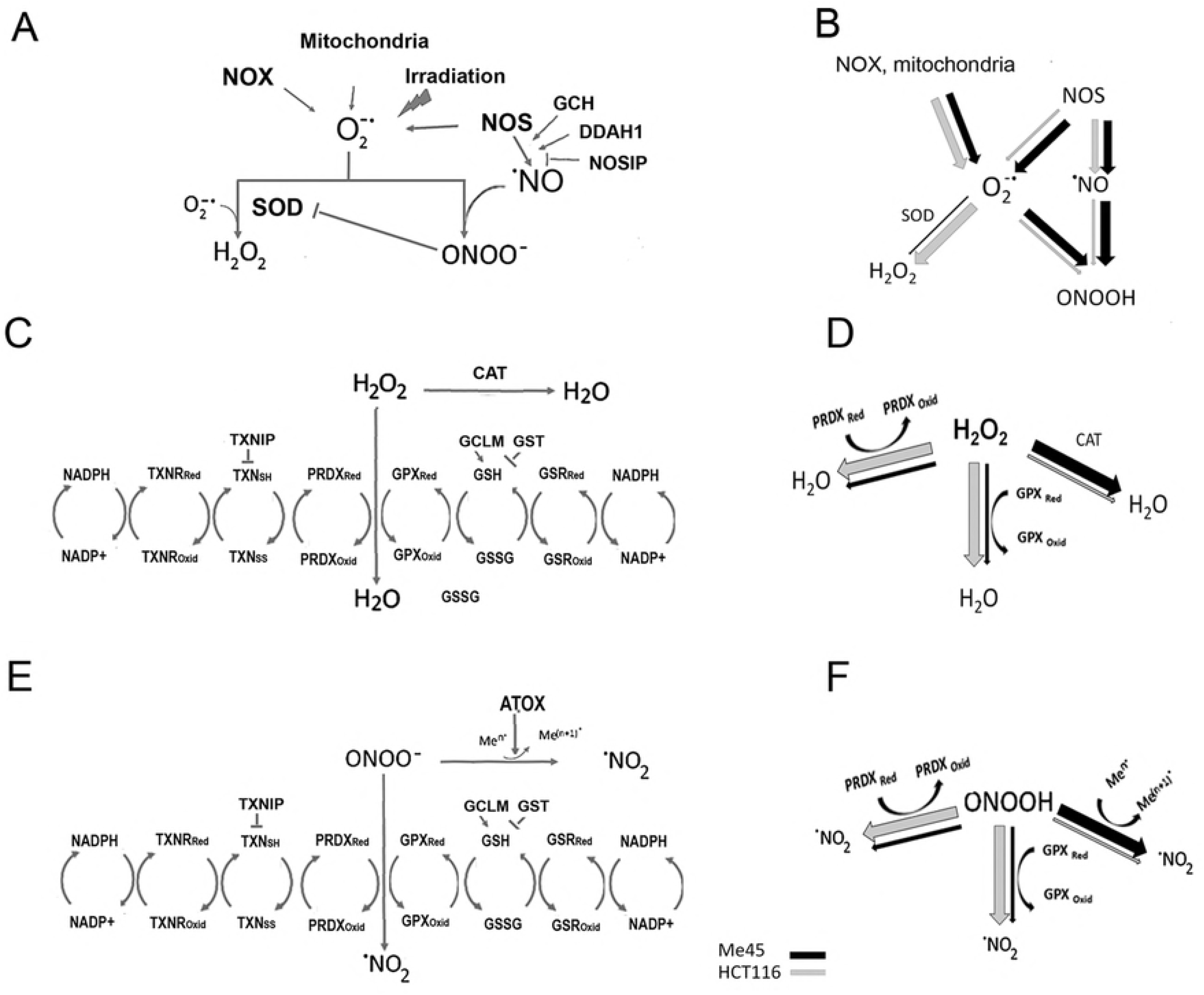
The main pathways for regulation of superoxide and NO levels in Me45 and HCT116 cells. A,C and E show production and further interactions of superoxide (A), hydrogen peroxide (C) and peroxynitrite (E) and regulatory pathways engaged. B,D and F compare use of presented regulatory pathways in HCT116 and Me45 cells by the size of black (Me45) and white (HCT116) arrows.

The two main pathways leading to regulation of superoxide levels start by its conversion to H_2_O_2_ or to peroxynitrite in reactions with NO (53,50). H_2_O_2_ may be created by interaction of two superoxide molecules, either spontaneously or more efficiently by superoxide dismutase (SOD) (53,24,22). Interaction of superoxide with NO starts another pathway by creation of the very reactive peroxynitrite radical (ONOO^-^); the sources of superoxide and NO and their spatial separation may determine further regulatory pathways through H_2_O_2_ or ONOO^-^ in cells.

NOS produces either NO or superoxide in appropriate conditions (54), and we speculate that this could explain the more frequent colocalization of these two types of radical in Me45 than in HCT116 cells (**Fig 4**). All three isoforms of NOS contain the N-terminal oxygenase and C-terminal reductase domains separated by a linker, and function as homodimers which produce NO by oxidation of L-arginine to L-citrulline (55 reviewed in 56,57). In the absence of the cofactor, tetrahydrobiopterin (BH4), the domains become uncoupled and NOS produces superoxide instead of NO (42,43,57 reviewed in 56). The levels of transcripts for the NOS isoforms are rather low and are similar in HCT116 and Me45 cells, except that for NOS2 which is slightly higher in HCT116 cells (Table 3 and Supplementary Material). However, the gene *GCH1* which encodes the rate-limiting enzyme in synthesis of BH4 (58) is expressed at a significantly lower level in Me45 cells (Table 3) which could result in insufficient availability of BH4 and consequently an increased production of superoxide by NOS. Further, in Me45 cells the level of transcripts for glutathione transferases is significantly higher (Table 2) and glutathionylation of NOS results in increased production of superoxide (59, 43,). Either or both of these scenarios would result in superoxide forming a larger fraction of the products of NOS in Me45 cells and to the observed more frequent apparent colocalization with NO. This would lead to higher production of peroxynitrite which may be further converted to NO_2_ by peroxiredoxins and glutathione peroxidases which also participate in reduction of H_2_O_2_ (60, 61) and these pathways are probably used preferentially by HCT116 cells which show higher expression of PRDX, TXN, GPX than Me45 cells. The other pathway for ONOO-reduction is interaction with transition metal centers (reviewed in 48) and Me45 cells show significantly higher levels than HCT116 cells of *ATOX* gene transcripts coding for copper chaperone (62,22) and of transcripts of thioredoxin-inhibiting protein TXNIP, suggesting that in Me45 cells interaction of ONOO^-^ with transition metals may be dominating.

Glutathione is a further important player in redox regulation, and its level is lower in Me45 cells than in HCT116 cells (Fig 5). This could plausibly be due to the lower expression of the *GCLM* gene (Table 3), or to greater use of glutathione for glutathionylation of proteins since genes coding for GSTs are more highly expressed in Me45 cells. As glutathione is necessary for reactivation of GPX, one could again expect that the pathway engaging GPX will be also less efficient in Me45 cells.

Redox balance plays a critical role in regulating biological processes and many cellular pathways, including stimulation and inhibition of proliferation, are influenced by ROS levels. Our results suggest that cells may concentrate on strict regulation of superoxide or hydrogen peroxide levels when changed by stress, and that stimulation or inhibition of cell proliferation depend on the dynamics of level fluctuations and less on the ROS levels themselves. We show for the first time that varying responses of different cell types to the same stimulus such as a specific dose of UVA may result from their use of different redox control pathways.

## Acknowledgments

This work was supported by the Polish National Science Center Grant 2015/19/B/ST7/02984.

Ronald Hancock (Laval University, Québec, Canada) is acknowledged for critically reading and editing the manuscript.

## Supplement

**Table S1**

**Expression of genes engaged in redox processes in Me45 and HCT116 cells**

